# Early maturation and hyperexcitability is a shared phenotype of cortical neurons derived from different ASD-associated mutations

**DOI:** 10.1101/2022.11.02.514882

**Authors:** Yara Hussein, Utkarsh Tripathi, Ashwani Choudhary, Ritu Nayak, David Peles, Idan Rosh, Tatiana Rabinski, Jose Djamus, Gad Vatine, Ronen Spiegel, Tali Garin-Shkolnik, Shani Stern

## Abstract

Autism Spectrum Disorder (ASD) is characterized mainly by social and sensory-motor abnormal and repetitive behavior patterns. Over hundreds of genes and thousands of genetic variants were reported to be highly penetrant and causative of ASD. Many of these mutations cause comorbidities such as epilepsy and intellectual disabilities (ID). In this study, we measured cortical neurons derived from induced pluripotent stem cells (iPSCs) of patients with four mutations in the genes *GRIN2B, SHANK3, UBTF*, as well as chromosomal duplication in the 7q11.23 region and compared them to neurons derived from a first-degree relative without the mutation. Using a whole-cell patch-clamp, we observed that the mutant cortical neurons demonstrated hyperexcitability and early maturation compared to control lines. These changes were characterized by increased sodium currents, increased amplitude and rate of excitatory postsynaptic currents (EPSCs), and more evoked action potentials in response to current stimulation in early-stage cell development (3-5 weeks post differentiation). These changes that appeared in all the different mutant lines, together with previously reported data, indicate that an early maturation and hyperexcitability may be a convergent phenotype of ASD cortical neurons.

## INTRODUCTION

Autism Spectrum Disorder (ASD) was first defined by Leo Kanner in 1943 and named “early infantile autism” as an independent disorder from the psychotic disorder of schizophrenia, describing 11 children with social, biological, and emotional abnormalities such as the inability to relate to others or objects in a traditional way, an anxious and obsessive desire for consistency, early eating difficulties and hearing problems^1^. Recently, more symptoms entered the category of autistic-like behaviors such as attention deficit hyperactivity disorder (ADHD), poorly integrated verbal and non-verbal communication, abnormalities in eye contact, hyper or hypo-reactivity to sensory input, repetitive body movements, and more^2,3^.

On the genomic aspect, there has been more attention to many variants in which chromosomal subregions are deleted or duplicated in an inherited and de novo manner as well^2^. Various genes and mutations have been reported to be associated with ASD^4^, and there seem to be a most definitive leaders: *ADNP, CHD8*, and *SHANK3*^5^.

On the neurobiological aspect, no specific brain area nor system has been confirmed to be entirely associated with the disorder, but an overall brain impairment has been shown starting from childhood^3^. The areas in the brain that are thought to be affected include cortical as well as non-cortical regions: the prefrontal cortex (PFC), Brodmann’s areas 9, 21, and 22, orbitofrontal cortex (OFC) fusiform gyrus, fronto-insular cortex, cingulate cortex, hippocampus, amygdala, cerebellum, and brainstem^6^. Our study focuses on the following four genetic mutations:

### Dup7

7q11.23 Duplication Syndrome, briefly known as **Dup7**, is caused by a duplication of 1.5-1.8 Mb in section q11.23 of chromosome 7^7^, also known as the Williams-Beuren syndrome critical region (WBSCR) and is inherited in an autosomal dominant manner^8^. Extra copies of the genes are located in the critical region and include the Elastin gene (ELN)-encoding elastin, a structural protein and a critical component of elastic fibers^9^, and the General Transcription Factor IIi gene (GTF2I)-encoding general transcription factor IIi, an inducible multifunctional transcription factor and a regulator of agonist-induced calcium entry to the cytoplasm^10^. These are thought to contribute significantly to the Syndrome’s symptoms^11,12^.

Common symptoms of Dup7 patients include anxiety disorders, selective mutism, ASD, ADHD, intellectual disability, seizures, severe speech delays, and hypotonia^13,14^ as well as Facial features including a broad forehead, a thin upper lip, and facial asymmetry^11^. Beyond that, magnetic resonance imaging (MRI) showed various abnormalities such as cerebral atrophy, increased cortical thickness, alterations in white matter’s volume and cerebellar hypoplasia, and an increased cortical extra-axial space^14^.

### SHANK3

The *SHANK* protein family, also known as ProSAP, including *SHANK 1-3*, is a family of scaffold proteins first identified using guanylate kinase-associated protein (GKAP) – a PSD-95-binding protein and a major component of the postsynaptic density (PSD), as bait.^15,16^ These proteins contain five key sites for protein-protein interaction: ankyrin repeats, Src homology 3 (SH3) domain, a vast proline-rich region, a C-terminal sterile α-motif (SAM) domain, and a PSD-95/discs large/zonula occludens-1 (PDZ) domain mediating its interaction with a variety of proteins as GKAP which moderates its binding to the NMDA and AMPA receptors^15,17,18^.

The cognate gene, ***SHANK3*** gene, is located at the terminal long arm of chromosome 22, coding for master scaffolding protein found in the body’s tissues and notably in the brain. It plays a critical role in the postsynaptic density of glutamatergic synapses and synaptic functions^15,19^. Mutations in this gene have been linked to deficits in synaptic function and plasticity, accompanied by lower reciprocal social interactions^18,20–23^, ASD, and other neurodevelopmental symptoms^21,24^. The individuals affected by the 22q13 deletion syndrome^24,25^, also known as Phelan-McDermid syndrome (PMS), suffer from developmental delays, a low muscular tone, a weakened perception of pain, delayed speech, seizures and ASD^26–28^. The patients also exhibit cortical alterations^29^.

### GRIN2B

Glutamate Ionotropic Receptor NMDA type Subunit 2B (***GRIN2B***) is a gene located on the ‘p’ arm of chromosome 12 (12p13.1 is its exact location)^30,31^. This gene is one member of a family of genes encoding various proteins that together form the NMDA receptor^32^ – glutamate-gated ion channel, which allows positively charged particles to flow through cells in the brain, activating neurons to send signals to each other^33,34^.

Mutations in this gene lead to a production of a nonfunctional GluN2B protein or completely prevent the production of GluN2B proteins from one copy of the gene in each cell; A shortage or dysfunction of this protein may cause an extreme reduction of the number of the functional NMDA receptors^33,35,36^, causing neurodevelopmental disorders^37^. This disorder is associated with West syndrome, characterized by intellectual disability, delayed development of speech and motor skills, focal seizures, weak muscle tone (hypotonia), pediatric encephalopathy, movement disorders, schizophrenia, and behavioral delays common in ASD^30,37–42^. In addition, MRI scans revealed a consistent malformation of cortical development (MCD) consistent with that of tubulinopathies, hypoplastic corpus callosum of varying degrees, enlarged and mildly dysplastic basal ganglia, hippocampal dysplasia as well as generalized cerebral volume loss^43^. *GRIN2B*-related disorders seem to have the highest prevalence among all GRIN-related disorders with 5.91 predicted incidence per 100,000 births^44^; This was previously reported to make up ∼ 0.22% of the total neurodevelopmental disorders^43^.

### UBTF

The upstream Binding Transcription Factor (***UBTF)*** is a gene that codes for Upstream Binding Factor (UBF) – a protein that acts as a transcription factor in RNA polymerase I (Pol I). UBF is critical for ribosomal RNA transcripts (rRNA) and synthesis from ribosomal DNA (rDNA) in the nucleolus^45^; A loss of UBF induces nuclear disruptions, including inhibition of cell proliferation, rapid and synchronous apoptosis as well as cell death^46^. Moreover, mutations in the *UBTF* gene lead to a production of an increased amount of ribosomal RNA transcript (rRNA), inducing depletion of RNA binding proteins, altered disposal machinery, and ribosome biogenesis^47^. These disruptions lead to pediatric neurodevelopmental regression starting at young onset (2.5 to 3 years). The degeneration starts with cognitive-motor deficits, reported in humans as well as knockout mice^48^. This regression is accompanied by dystonia, parkinsonism, severe intellectual disability, feeding difficulties, autistic-like behaviors, a slow loss of motor, cognitive, and speech capabilities, as well as severe epilepsy ^47,49–52^. Recent MRI studies showed gradual widespread brain atrophy in both supratentorial and infratentorial areas affecting both grey and white matter in patients with similar neuroregression patterns^47,50^. Furthermore, increased pre-rRNA and 18S rRNA expression was reported in addition to nucleolar abnormalities quantified by quantitative real-time polymerase chain reaction (qRT-PCR), increased number of DNA breaks, defective cell cycle progression by comet assay, and apoptosis by TUNEL assay^47^.

## MATERIALS AND METHODS

After obtaining institutional committee approval and written informed consent from all participants or their respective legal guardians, iPSCs were generated from the peripheral blood mononuclear cells (PBMCs) of one male child carrying *Dup7* (7q11.23 dup) (see supplementary Fig S1a), one female child carrying *GRIN2B* (c.2065 G->T), one female child carrying *SHANK3*^53^ (C.3679insG) (see supplementary Fig S1b)and one female child carrying *UBTF* (E210K) mutations and a corresponding number of controls sharing the same genetic backgrounds (first-degree relatives from the same gender, except for *GRIN2B*), Table S1 presents a description of the patients’ cohort.

### PBMC isolation

Participants had undergone a regular blood test (except for the *SHANK3* cohort and *GRIN2B*-mutant line, in which iPSCs were generated from skin biopsies); blood was collected into heparin-coated tubes, diluted with phosphate-buffered saline (PBS) and centrifuged for 30 minutes at 1800□×g at 23□C. The mononuclear cells (MCs) were collected by pipetting the buffy coat – the cell layer between the gel barrier and the plasma, into a sterile 15 ml conical centrifuge tube, adding PBS to reach a volume of 10 ml. This was followed by centrifuging (300 xg at room temperature for 15 min), resuspended using fetal bovine serum (FBS, F7524 Merck) 10% DMSO to freeze a total of ∼2×10^6^ cells per cryovial in 1ml volume were aliquoted, frozen and kept in liquid nitrogen.

### Reprograming of PBMCs into iPSCs

PBMCs were grown in a 24-well plate and seeded in fresh StemPro™ SFM medium (10639011, Thermo Fisher Scientific). Sendai viral particle factors from CytoTune-iPS Sendai Reprogramming Kit (A16518, Life Technologies, Carlsbad, CA, USA) were added based on the manufacturer’s recommendations^54^, followed by centrifuging the cells (1,160□×g, at 35°C, for 30 minutes) and seeded on a 24-well plate and an overnight incubation in 5% CO_2_ at 37°C. The next day, the cells were transferred into a low attachment coated 12-well plate. Later, they were plated in six-well Matrigel-coated plates for three days and fed with a complete PBMC medium: StemPro™-34 supplement and2mM of L-Glutamine, followed by two days of gradually transitioning into iPSC medium (DMEM/F-12, GlutaMAX, KnockOut™ Serum Replacement, 10mM MEM Non-Essential Amino Acids Solution and 55mM β-mercaptoethanol). This was followed by daily replacing spent medium with fresh mTeSR for 20 days. When iPSC colonies were large enough, manual picking of the colonies was done, and colonies were transferred onto six well-Matrigel-coated plates. Immunocytochemistry of TRA-1-60, SOX2, and OCT4 antibodies confirmed pluripotency.

### Generating neuronal cultures

Cortical neurons were plated based on a previously described protocol^55^. In brief, iPSCs were grown to ∼80% confluency, embryonic bodies (EBs) were formed by mechanical dissociation using dispase for 20 minutes and plated onto low-adherence plates in mTeSR medium with ROCK inhibitor for one day, followed by one day of mTeSR medium only. On the following ten days, cells were fed with EB media: (DMEM/F12 with Glutamax(1:100), B27 with Retinoic Acid (RA) (1:100), N2 supplement (1:100) and 0.1uM LDN), followed by plating onto polyornithine/laminin (Sigma)-coated dishes in DMEM/F12 (Invitrogen) plus N2, B27 and laminin for the following seven days. The rosettes were selected based on their morphology and were manually picked, dissociated with Accutase (Chemicon) and plated onto poly-l-ornithine/laminin-coated plates and fed with complete NPC medium (DMEM/F12 with Glutamax (1:100), B27 supplement with RA (1:100), N2 supplement (1:100), laminin (1 mg/ml) and 20ng/ml FGF2) for the following 15-20 days (based on their confluency)… NPCs were then differentiated into cortical neurons by feeding with the differentiation medium that contained: DMEM/F12, N2, B27, Glutamax, ascorbic acid (200 nM), cyclic AMP (cAMP; 500 mg/ml), laminin (1 mg/ml), BDNF (20 ng/ml), GDNF (20 ng/ml) for ten days. Between days 11-14, the cells were dissociated again and then fed with Brainphys medium with B27 supplement (1:100), N2 supplement (1:100), ascorbic acid (200 nM), cyclic AMP (500 mg/ml), BDNF (20 ng/ml), GDNF (20 ng/ml) and laminin (1 mg/ml).

### Electrophysiology

Whole-cell patch-clamp recordings have been performed on neurons derived from patients carrying the *Dup7, GRIN2B, SHANK3*, and *UBTF* mutations, as well as neurons derived from healthy controls based on previously described^56^ with some modifications^57^, three to five weeks post differentiation. Culture coverslips were placed inside a recording chamber filled with HEPES-based artificial cerebrospinal fluid (ACSF) containing (in mM): 10 HEPES,139 NaCl, 4 KCl, 2 CaCl_2_, 10 D-glucose, and 1MgCl_2_(pH 7.5, osmolarity adjusted to 310 mOsm) at room temperature. The recording micropipettes (tip resistance of 9-12 MΩ) were filled with an internal solution containing (in mM): 130 K-gluconate, 6 KCl, 4 NaCl, 10 Na-HEPES, 0.2 K-EGTA, 0.3 GTP, 2 Mg-ATP, 0.2 cAMP, 10 D-glucose, 0.15% biocytin and 0.06% rhodamine (pH 7.5, osmolarity adjusted to 290-300 mOsm). All measurements were done at room temperature using Clampex v11.1 with a sampling rate of 20 kHz.

### Analysis of electrophysiological recordings

Analysis was performed based on previously described^57^ analysis using custom-written MATLAB scripts modified as follows:

#### Synaptic currents analysis

The mean and standard error (SE) of the excitatory postsynaptic currents (EPSCs) amplitudes for each active cell were calculated. The cumulative distribution of EPSCs amplitude was calculated for each group. For each cell, the rate of the events was calculated by dividing the number of events by the time period of the recording (non-active cells were included and had an event rate=0). The mean of all cells’ rates and the standard error of the frequencies were computed for the control and mutant groups. Non-parametric statistical tests (Wilcoxon signed rank test) were performed for comparisons.

#### Sodium, fast, and slow potassium currents

Neurons were held in voltage clamp mode at -60 mV, and voltage steps of 400 ms were performed in the -100 to 90 mV range. Currents were typically normalized by the cells’ capacitance; The sodium current was computed by subtracting the sodium current after stabilization from the lowest value of the inward sodium current. The fast potassium currents were measured by the maximum outward currents that appeared within a few milliseconds after a depolarization step. The slow potassium currents were measured after the 400 ms depolarization phase. A one-way ANOVA test was performed for the statistical analysis.

#### Evoked action potentials (APs)

Neurons were held in current clamp mode at -60 mV with a constant holding current. Following this, current injections were given in 3-pA steps throughout 400 ms, starting 12 pA below the steady-hold current. A total of 38 depolarization steps were given. The total evoked action potential was the total number of action potentials that were counted in the 38 depolarization steps. Non-parametric statistical tests (Wilcoxon signed rank test) were performed for comparisons.

#### Action potential shape analysis

The first evoked action potential generated with the lowest amount of injected current was used for spike shape analysis. The spike threshold was defined as the membrane potential at which the slope of the depolarizing membrane potential increased dramatically, resulting in an AP (the second derivative of the voltage versus time as the initial maximum). The spike height was calculated as the difference between the highest membrane potential during a spike and the threshold. The spike rise time is the time it takes the spike to reach the maximum. The spike width was calculated as the time it took the membrane potential to reach half of the spike amplitude in the rising part.

### Analysis of brightfield images

Brightfield images of NPCs from both control and patient cultures were captured using a Nikon Eclipse Ts2 microscope with 20X magnification. Neurite’s length was measured manually using ImageJ and statistically tested with Wilcoxon’s signed rank test and visualized using MATLAB.

### Immunocytochemistry (ICC)

Cells were fixed in 4% paraformaldehyde for 15 minutes, followed by three washes of DPBS, blocked and permeabilized in PBS containing 0.1%-0.2% Triton X-100 and 10% horse serum. The coverslips were incubated with primary antibodies; for NPCs: rabbit anti-PAX6 (CST, mAb#60433, 1:250) and mouse-anti NESTIN (CST, mAb#33475,1:2000); for Neurons: chicken-anti MAP2 (abcam, ab92434, 1:500) and rabbit anti-TBR1 (abcam, ab183032, 1:250) and Rabbit-anti VGLuT1 (abcam, ab227805, 1:500) and Mouse-anti GABA (abcam, ab86186, 1:400) in the blocking solution overnight at 4□C. On the next day, they were washed in DPBS and incubated with DAPI (abcam, ab228549, 1:1000) and corresponding secondary antibodies (abcam, ab150084, ab150117, ab175711, 1:250) for 60 minutes at room temperature. Then the coverslips were washed three times, mounted on glass slides using Fluromount-G (mounting medium), and dried overnight while being protected from light. Fluorescence signals were detected using a Leica THUNDER imager and analyzed using ImageJ and MATLAB

## Code Availability

The custom-written scripts used for the analysis can be shared upon reasonable request.

## RESULTS

### Dup7 cortical neurons display increased sodium and potassium currents, increased synaptic activity and hyperexcitability early in the differentiation

We performed whole-cell patch clamp experiments five weeks (day 34) after the start of the differentiation of 16 Dup7-mutant neurons and 13 control neurons derived from a first-degree relative of the same gender. In a voltage clamp mode, EPSC recordings were performed by holding the cell at -60mV. We observed an increase in the rate of EPSCs of the mutant neurons compared to the controls (0.13 ± 0.08 Hz in Dup7-mutant neurons and 0.07 ± 0.07 Hz in the control neurons, p=0.03), as shown in Fig. 1a-c. Fig. 1a-b presents representative traces, while Fig. 1c represents the average over all the recordings. Additionally, a significant increase in the mean amplitude of the EPSCs was observed. The Dup7-mutant neurons had a larger amplitude compared to the control neurons (12.72 ± 2.86 pA for the mutant neurons and 7.12 + 5.09 pA for the control neurons (p=0.002, (Fig. 1d.)). The cumulative distribution of the EPSC amplitudes for Dup7-mutant neurons is slightly right-shifted compared to control neurons indicating larger amplitudes of EPSCs (Fig. 1e).

**Fig. 1.**
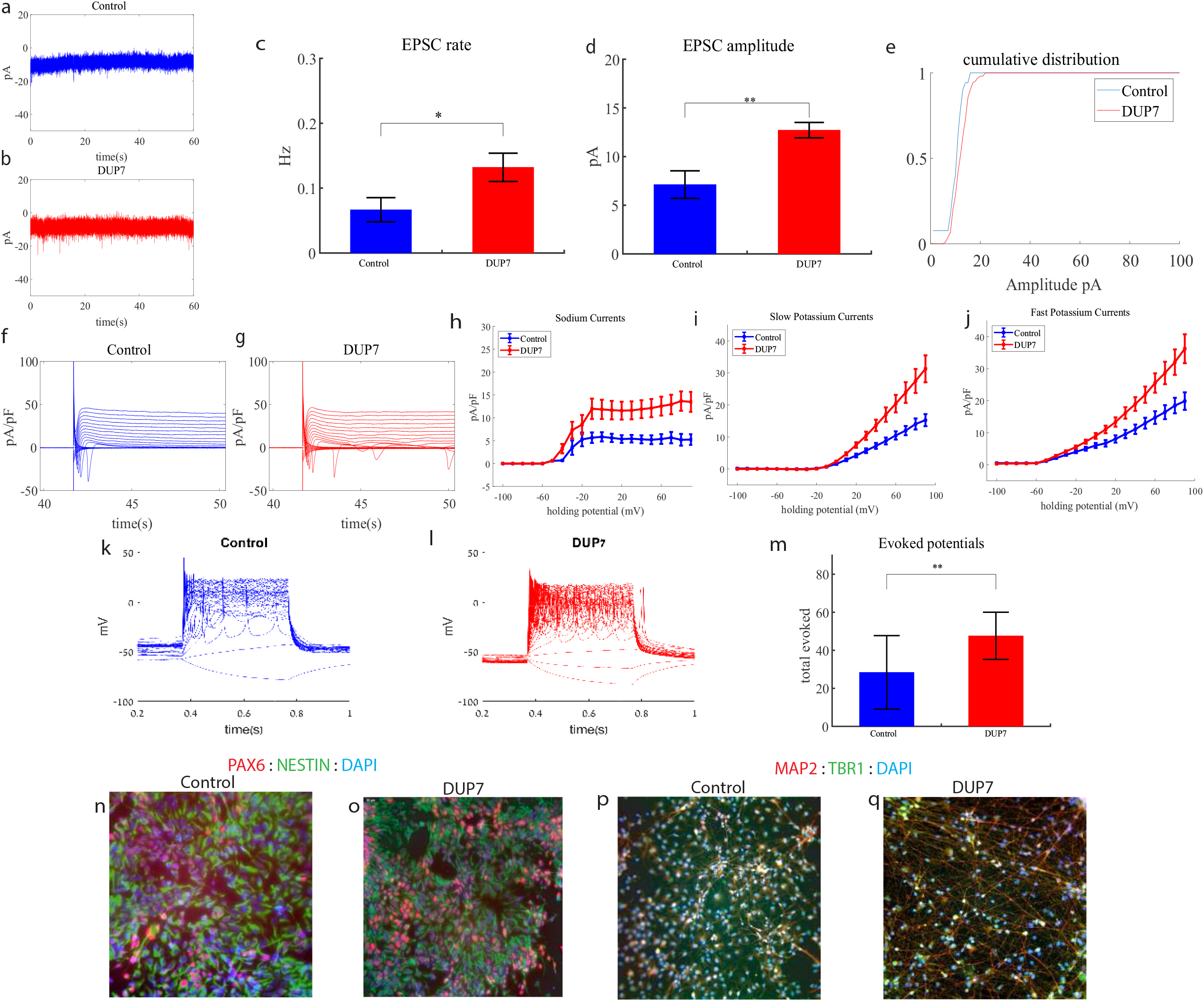
Young (5 weeks post differentiation) Dup7-mutant neurons are hyperexcitable compared to control neurons. **a** A representative trace of (EPSCsmeasured in control cortical neurons at five weeks post-differentiation. **b** A representative trace of EPSCs measured in a dup7-mutant neuron at five weeks post-differentiation. **c** The mean rate of synaptic events was higher in the dup7-mutant neurons compared to control neurons (p=0.029). **d** The average amplitude of EPSCs was increased in the dup7-mutant neurons (p =0.002). **e** The cumulative distribution of the amplitude of EPSCs is slightly right-shifted in the dup7-mutant neurons, indicating an increase in the amplitudes. **f** A Representative trace of sodium and potassium currents recorded in a voltage-clamp mode in control neurons. **g** A Representative trace of sodium and potassium currents recorded in a voltage-clamp mode in dup7-mutant neurons. **h** The average sodium currents in dup7-mutant neurons is increased compared to control neurons (p=0.004). **i** The average slow potassium currents in dup7-mutant neurons is increased compared to control neurons (p=0.03). **j** The average fast potassium currents is increased in dup7-mutant compared to control neurons (p=0.01). **k** A representative recording of evoked action potentials in a current-clamp mode of a control neuron. **l** A representative recording of evoked action potentials in a current-clamp mode of a dup7-mutant neuron. **m** The total number of evoked action potentials is larger in dup7-mutant neurons compared to control neurons (p=0.006). **n**,**o** A representative image of control **n** and mutant **o** NPCs that were immunostained for DAPI, PAX6, and Nestin. **p-q** A representative image of control **p** and mutant **q** neurons that were immunostained for DAPI, MAP2, and TBR1.

Next, we recorded in voltage clamp mode the sodium and potassium currents. We observed a significantly larger normalized sodium current in the Dup7-mutant neurons compared to the control neurons (F (1,38) = 9.43, p=0.004). Representative traces of the recordings are shown in Fig. 1f (control) and 1g (mutant). The average sodium currents are presented in Fig. 1h. Additionally, we observed increased slow and fast potassium currents (normalized by the capacitance) in the Dup7-mutant neurons compared to controls (F (1,14) = 5.61; p=0.03 for the slow potassium currents and F (1,14) = 8.36; p=0.01 for the fast potassium currents) over the 10-80 mV range (Fig. 1h-j). We next measured the number of evoked action potentials in a current clamp mode as a measure of the neuronal excitability. We observed a hyperexcitability pattern for the Dup7-mutant neurons compared to control neurons. The total number of evoked potentials (see Methods) for the Dup7-mutant neurons was 47.63 ± 49.61; for the control neurons, it was 28.5 ± 69.48 (p=0.006). A representative example is presented in Fig. 1k (control) and 1l (Dup7), and the average over all recordings is shown in Fig. 1m. Spike shape analysis (see Methods) is presented in Table S2 .Examples of ICC images for control and mutant lines are shown in Fig. 1n-o. (typical NPC markers PAX6 and NESTIN) and 1p-q. (neuronal markers MAP2 and the cortical marker TBR1).

### *GRIN2B* cortical neurons display increased sodium and potassium currents and hyperexcitability early in the differentiation

We performed whole-cell patch clamp experiments three weeks (day 19) after the start of the differentiation of 12 *GRIN2B*-mutant neurons and five weeks (day 27) of 7 control neurons. In a voltage clamp mode, EPSC recordings were performed by holding the cell at -60mV. We observed an increase in the rate of EPSCs of the mutant neurons compared to the controls (0.39± 0.27 Hz in *GRIN2B*-mutant neurons and 0.17 ± 0.2 Hz in the control neurons, p=0.03) as shown in Fig. 2a-c. Fig. 2a-b presents representative traces, while Fig. 2c. represents the average over all the recordings. No significant difference in the mean amplitude of the EPSCs was observed (Fig. 2d.). The cumulative distribution of the EPSC amplitudes was similar between the control and *GRIN2B*-mutant neurons (Fig. 2e.).

**Fig. 2.**
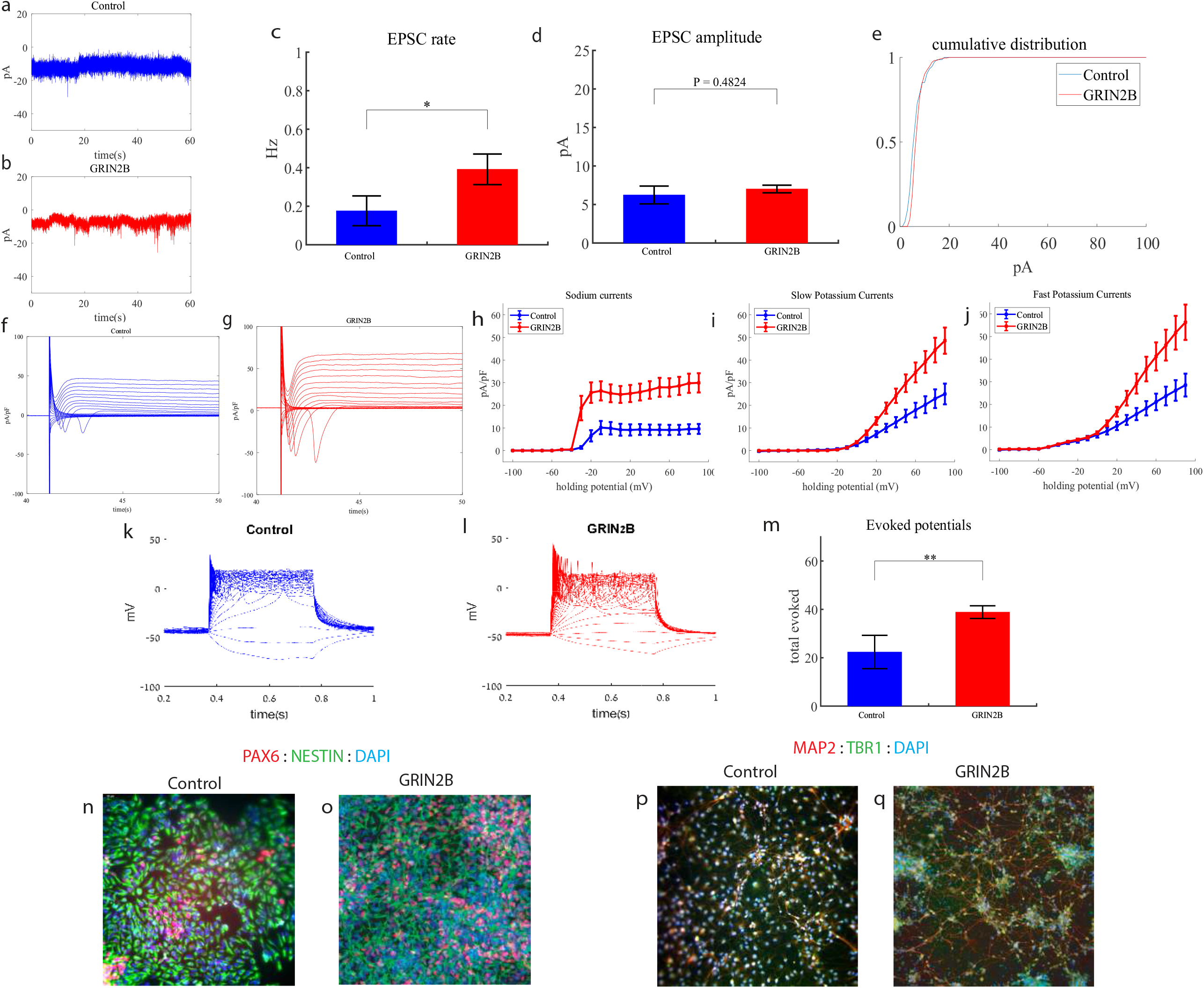
Young (3 weeks post differentiation) *GRIN2B*-mutant neurons are hyperexcitable compared to control neurons (5 weeks post differentiation). **a** A representative trace of EPSCs measured in control cortical neurons at five weeks post-differentiation. **b** A representative trace of EPSCs measured in a *GRIN2B*-mutant neuron at three weeks post-differentiation. **c** The mean rate of synaptic events was higher in the *GRIN2B*-mutant neurons (p=0.03). **d** The mean amplitude of EPSCs was increased but not significantly different in the *GRIN2B*-mutant neurons. **e** The cumulative distribution of the amplitude of the EPSCs of *GRIN2B*-mutant and control neurons looks similar. **F** A Representative trace of sodium and potassium currents recorded in a voltage-clamp mode in control neurons. **g** A Representative trace of sodium and potassium currents recorded in voltage-clamp in *GRIN2B*-mutant neurons. **h** The average sodium currents in *GRIN2B*-mutant neurons is severely increased compared to control neurons (p=0.006). **I** The average slow potassium currents in *GRIN2B*-mutant neurons is increased compared to control neurons (p=0.04). **j** The average fast potassium currents in *GRIN2B*-mutant neurons is increased compared to control neurons (p=0.03). **k** A representative recording of evoked action potentials in a current-clamp mode of a control neuron. **l** A representative recording of evoked action potentials in a current-clamp mode of a *GRIN2B*-mutant neuron. **m** The total number of evoked action potentials is larger in the *GRIN2B*-mutant neurons compared to control neurons (p=0.003). **n**,**o** A representative image of control (**n**) and mutant (**o**) NPCs that were immunostained for DAPI, PAX6, and Nestin. **p-q** A representative image of control (**p**) and mutant (**q**) neurons that were immunostained for DAPI, MAP2, and TBR1.

Next, we recorded in voltage clamp mode the sodium and potassium currents. We observed a significantly larger normalized sodium current in the *GRIN2B*-mutant neurons compared to the control neurons (F (1,38) = 14.1, p=0.0006). Representative traces of the recordings are shown in Fig. 2f. (control) and 2g. (mutant). The average sodium currents are presented in Fig. 2h. Additionally, we observed larger slow and fast potassium currents (normalized by the capacitance) in the *GRIN2B*-mutant neurons compared to controls (F (1,16) = 4.6, p=0.04 for the slow potassium currents and F (1,16) = 5.39, p = 0.03 for the fast potassium currents) over the 0-80 mV range (Fig. 2h-j).

We next measured the number of evoked action potentials in current clamp mode as a measure of the neuronal excitability. We observed a hyperexcitability pattern for the *GRIN2B*-mutant neurons compared to control neurons. The total number of evoked potentials (see Methods) in the *GRIN2B*-mutant neurons was 38.86 ± 10.17, and in the control neurons, it was 22.37 ± 19.5 (p=0.003). A representative example is presented in Fig. 2k (control) and 2l (*GRIN2B*), and the average over all recordings is presented in Fig. 2m. Furthermore, we observed a significant increase in *GRIN2B*-mutant neurons’ spike amplitude compared to control neurons (41.2 ± 12.5 mV in *GRIN2B*-mutant neurons and 20.5 ± 14.01 mV in control neurons, p =0.008); further spike shape analysis is presented in Table S2. Examples of ICC images for control and mutant lines are shown in Fig. 2n-o. (typical NPC markers PAX6 and NESTIN) and 2p-q. (neuronal markers MAP2, the cortical marker TBR1).

### *SHANK3* cortical neurons display increased sodium and slow potassium currents, a drastic increase in synaptic activity and hyperexcitability early in the differentiation

We performed whole-cell patch clamp experiments five weeks (days 29-32) after the start of the differentiation of 21 *SHANK3*-mutant and 17 control neurons. In a voltage clamp mode, EPSC recordings were performed by holding the cell at -60mV. We observed a significant increase in the rate of EPSCs of the mutant neurons compared to the controls (0.28 ± 0.36 Hz in *SHANK3*-mutant neurons and 0.08 ± 0.06 Hz in the control neurons, p0.003) as shown in Fig. 3a-c. Fig. 3a-b presents representative traces, while Fig. 3c represents the average over all the recordings. Additionally, a drastic increase in the mean amplitude of the EPSCs was observed. The *SHANK3*-mutant neurons had a larger amplitude compared to the control neurons (10.145 ± 3.26 pA for the mutant neurons and 5.29 + 1.65 pA for the control neurons, p=1.15e^-5^, (Fig. 3d)). The cumulative distribution of the EPSC amplitudes for *SHANK3*-mutant neurons is right shifted compared to control neurons indicating larger amplitudes of EPSCs (Fig. 3e).

**Fig. 3.**
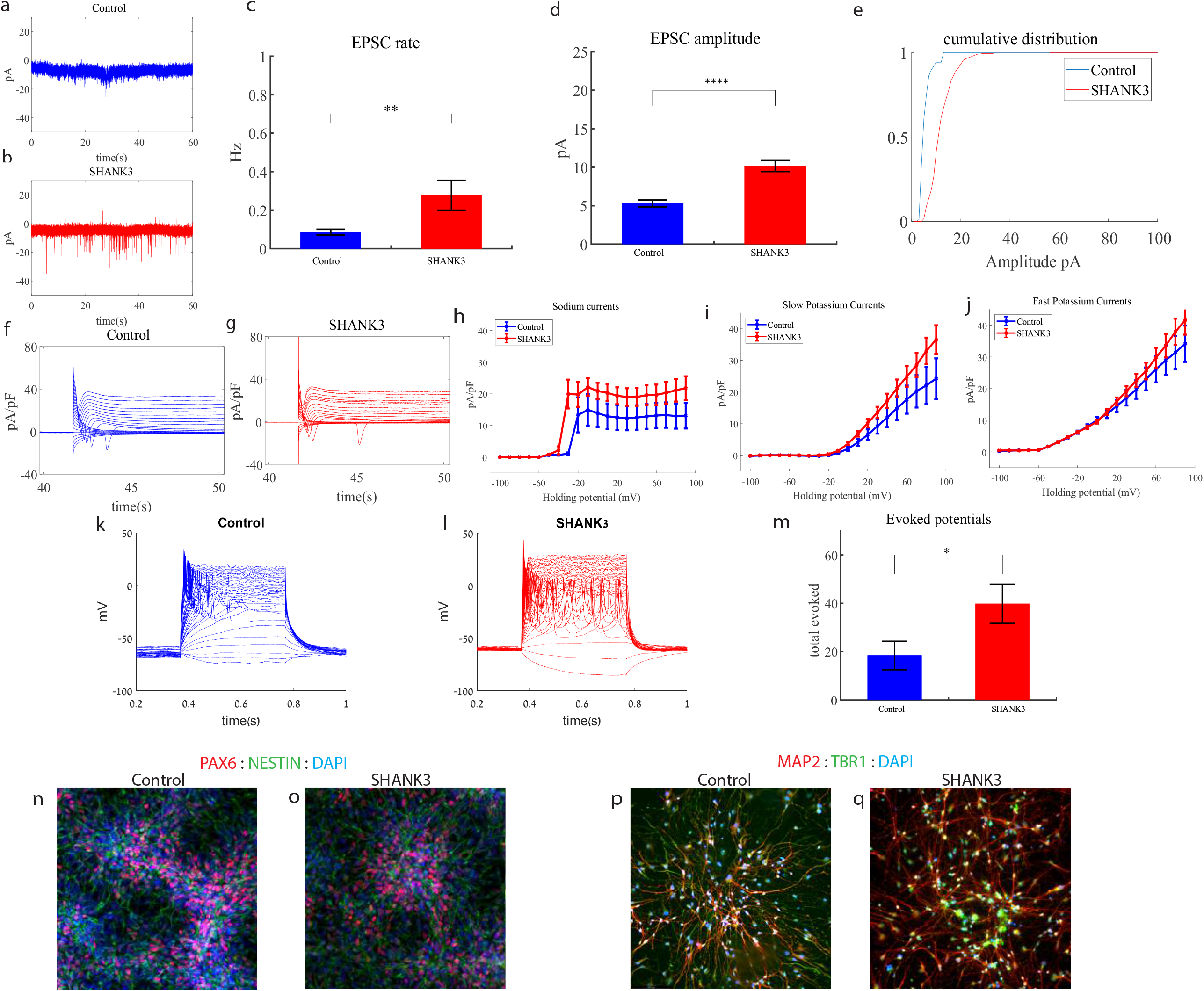
Young (5 weeks post differentiation) *SHANK3*-mutant neurons are hyperexcitable compared to control neurons. **a** A representative trace of excitatory postsynaptic currents (EPSCs) that were measured in control cortical neurons at five weeks post-differentiation. **b** A representative trace of EPSCs measured in a *SHANK3*-mutant neuron at five weeks post-differentiation. **c** The mean rate of synaptic events was higher in the *SHANK3*-mutant neurons (p=0.003). **d** The average amplitude of EPSCs was increased in the *SHANK3*-mutant neurons (p=1.15e^-5^). **e** The cumulative distribution of the amplitude of EPSCs of *SHANK3*-mutant is right-shifted, indicating an increase in the amplitudes. **f** A Representative trace of sodium and potassium currents recorded in voltage-clamp in control neurons. **g** A Representative trace of sodium and potassium currents recorded in a voltage-clamp mode in *SHANK3*-mutant neurons. **h** The average sodium currents in *SHANK3*-mutant neurons is increased compared to control neurons (p=0.04). **I** The average slow potassium currents in *SHANK3*-mutant neurons is increased compared to control neurons (p=0.03). **j** The average fast potassium currents look similar and not significantly different in *SHANK3*-mutant compared to control neurons. **k** A representative recording of evoked action potentials in a current-clamp mode of control neurons. **l** A representative recording of evoked action potentials in a current-clamp mode of *SHANK3*-mutant neurons. **m** The total number of evoked action potentials is larger in *SHANK3*-mutant neurons compared to control neurons (p=0.03). **n**,**o** A representative image of control (**n**) and mutant (**o**) NPCs that were immunostained for DAPI, PAX6, and Nestin. **p-q** A representative image of control (**p**) and mutant (**q**) neurons that were immunostained for DAPI, MAP2, and TBR1.

Next, we recorded in voltage clamp mode the sodium and potassium currents. We observed a significantly larger normalized sodium current in the *SHANK3*-mutant neurons compared to the control neurons F (1,36) = 4.51, p=0.04. Representative traces of the recordings are shown in Fig. 3f (control) and 3g (mutant). The average sodium currents are presented in Fig. 3h. Additionally, we observed increased slow, but not fast, potassium currents (normalized by the capacitance) in the *SHANK3*-mutant neurons compared to controls, (F (1,10) = 5.68; p=0.03) over the 40-90 mV range (Fig. 3h-j).

We next measured the number of evoked action potentials in current clamp mode as a measure of the neuronal excitability. We observed a hyperexcitability pattern for the *SHANK3*-mutant neurons compared to control neurons. The total number of evoked potentials (see Method) for the *SHANK3*-mutant neurons was 31.32 ± 39.8; for the control neurons, it was 18.4 ± 22.96 (p=0.03). A representative example is presented in Fig. 3k. (control) and 3l. (*SHANK3*) and the average over all recordings are presented in Fig. 3m. Furthermore, we observed a more depolarized threshold (higher) in the *SHANK3*-mutant neurons compared to control neurons (30.4 ± 5.6 mV in the *SHANK3*-mutant neurons and 25.5 ± 4.2 mV in control neurons, p =0.02); further spike shape analysis is presented in Table S2. Examples of ICC images of control and mutant lines are shown in Fig. 3n-o. (typical NPC markers PAX6 and NESTIN) and 3p-q. (neuronal markers MAP2, the cortical marker TBR1).

### *UBTF* cortical neurons display increased sodium currents, an increase in synaptic amplitude, an increase in spontaneous activity and hyperexcitability early in the differentiation

We performed whole-cell patch clamp experiments five to six weeks (days 32-37) after the start of the differentiation of 22 *UBTF*-mutant neurons and five to six weeks (days 32-37) after the beginning of the differentiation of 18 control neurons. In a voltage clamp mode, EPSC recordings were performed by holding the cell at -60mV. Fig. 4a-b presents representative traces, while Fig. 4c. represents the average EPSC amplitude over all the recordings. The EPSC rate of the *UBTF*-mutant neurons is slightly similar to the control neurons (Fig. 4c). We observed an increase in the mean amplitude of the EPSCs. The *UBTF*-mutant neurons had a larger amplitude compared to control neurons it was7.81± 1.16 pA for the mutant neurons and 6.55 + 2.39 pA for control neurons, p=0.02 (Fig. 4d.). The cumulative distribution of the EPSC amplitudes for *UBTF*-mutant neurons is slightly right-shifted compared to control neurons indicating larger amplitudes of EPSCs (Fig. 4e.).

**Fig. 4.**
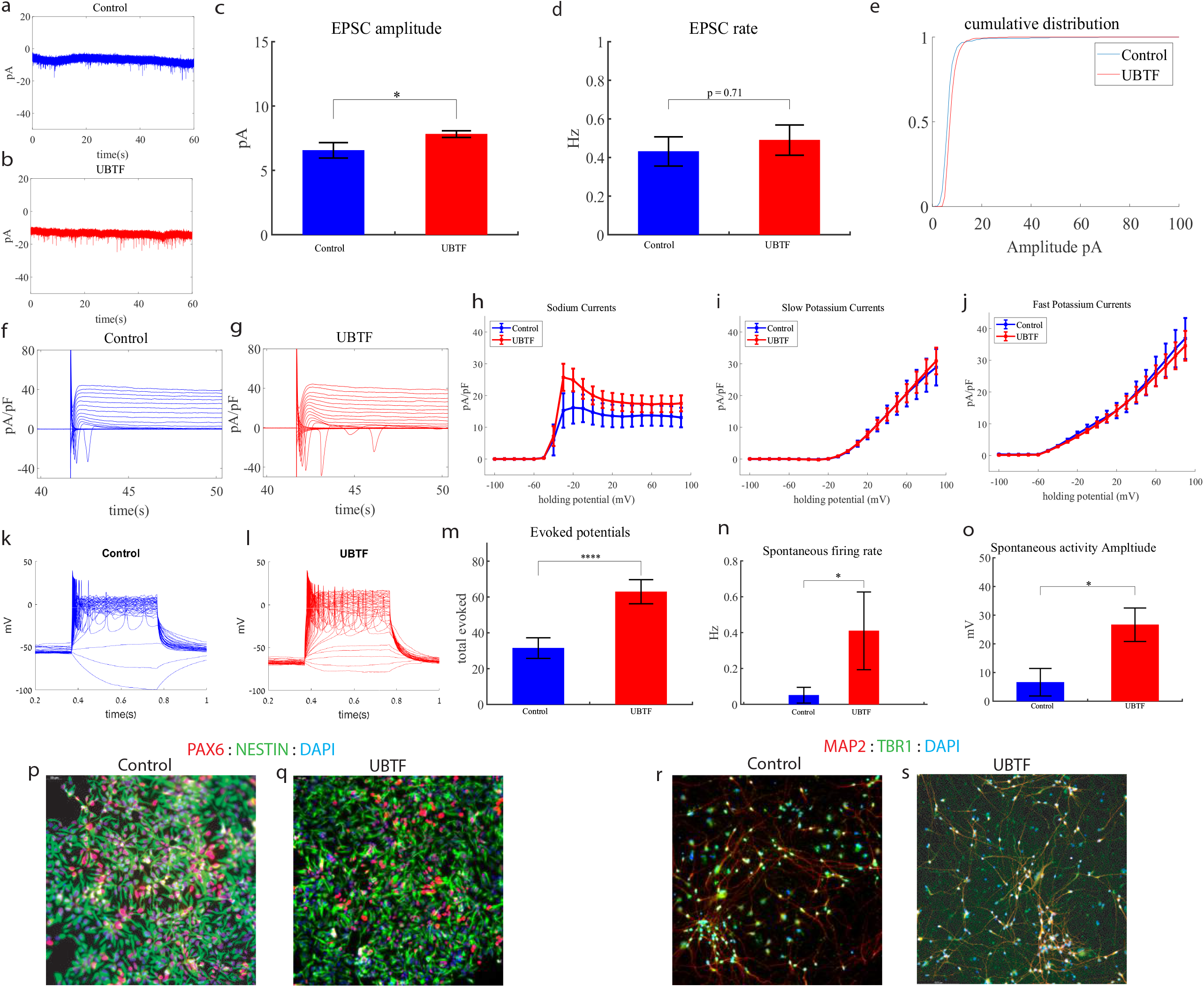
Young (5 weeks post differentiation) *UBTF*-mutant neurons are hyperexcitable compared to control neurons (6 weeks post differentiation). **a** A representative trace of excitatory postsynaptic currents (EPSCs) that were measured in control cortical neurons at six weeks post-differentiation. **b** A representative trace of EPSCs measured in a *UBTF*-mutant neuron at five weeks post-differentiation. **c** The mean rate of synaptic events was not significantly different in *UBTF*-mutant neurons compared to control neurons. **d** The average amplitude of EPSCs was increased in the *UBTF*-mutant neurons (p =0.02). **e** The cumulative distribution of the amplitude of EPSCs of *UBTF*-mutant neurons is slightly right-shifted, indicating an increase in the amplitudes. **f** A Representative trace of sodium and potassium currents recorded in a voltage-clamp mode in control neurons. **g** A Representative trace of sodium and potassium currents recorded in a voltage-clamp mode in *UBTF*-mutant neurons. **h** The average sodium currents in *UBTF*-mutant neurons is increased compared to control neurons (p<0.05). **I** The average slow potassium currents look similar in the *UBTF*-mutant and control neurons. **j** The average fast potassium currents look identical in the *UBTF*-mutant and control neurons. **k** A representative recording of evoked action potentials in the current-clamp mode of control neurons. **l** A representative recording of evoked action potentials in a current-clamp mode of *UBTF*-mutant neurons. **m** The total number of evoked action potentials is higher in the *UBTF*-mutant neurons compared to control neurons (p=5.56e^-4^). **n** The mean rate of spontaneous action potentials was significantly higher in UBTF-mutant neurons compared to control neurons (p = 0.013). **o** The mean amplitude of spontaneous action potentials in *UBTF*-mutant neurons was larger compared to control neurons (p = 0.012). **p**,**q** A representative image of control (**p**) and mutant (**q**) NPCs that were immunostained for DAPI, PAX6, and Nestin. **r-s** A representative image of control (**r**) and mutant (**s**) neurons that were immunostained for DAPI, MAP2, and TBR1.

Next, we recorded in voltage clamp mode the sodium and potassium currents. We observed a significantly larger normalized sodium current in the *UBTF*-mutant neurons compared to the control neurons F (1,28) = 5.58, p=0.03 over the -50-90-mV range. Representative traces of the recordings are shown in Fig. 4f. (control) and 4g. (mutant). The average sodium currents is presented in Fig. 4h. An ANOVA test indicated no significant differences in the slow and fast potassium currents (Fig. 4h-j.).

We next measured the number of evoked action potentials (see Methods). For the *UBTF*-mutant neurons, it was 62.95 ± 31.56 and for the control neurons, it was 31.46± 22.3 (p=5.56e^-4^). A representative example is presented in Fig. 4k. (control) and 4l. (*UBTF*) and the average over all recordings are presented in Fig. 4m. Furthermore, we observed a significant increase in *UBTF*-mutant neurons’ spike amplitude compared to control neurons (50± 17.05 mV in the *UBTF*-mutant neurons and 36.04 ± 18.01 mV in control neurons, p =0.03). Besides, we observed a narrower spike in the *UBTF*-mutant neurons compared to the control neurons (3.3 ± 1.5 ms in the *UBTF*-mutant neurons and 12.5 ±17.8 ms in the control neurons, p = 0.003); further spike shape analysis is presented in Table S2. The spontaneous neuronal activity (spontaneous action potentials) was measured in a holding potential of -45mV. We observed a significant increase in the spontaneous activity rate (p= 0.013) and amplitudes (p= 0.012) in the *UBTF*-mutant neurons compared to control neurons (Fig. 4n-o.). Examples of ICC images of control and mutant lines are shown in Fig. 4p-q. (typical NPC markers PAX6 and NESTIN) and 4r-s. (neuronal markers MAP2, the cortical marker TBR1).

### Mutant Neural Progenitor Cells Exhibit Longer Neurite Lengths Compared to Controls, alongside with decreased GABA-Positive Neurons in ASD-Related Mutant neuronal cultures

We observed that the lengths of neurites of neural progenitor cells (NPCs) in mutant groups were significantly increased compared to control groups (Fig. 5a). This difference was determined by analyzing brightfield images captured at 20X magnification and measuring the length of neurites using a standardized methodology (see Methods). Specifically, the average length of NPC neurites in the Dup7-mutant NPCs (left) was 0.12 ± 0.02 cm compared to control NPCs was 0.07 ± 0.02 cm (p=4.36e-04), in the *SHANK3*-mutant NPCs (middle) the average length was 0.09 ± 0.03 cm compared to control NPCs 0.07 ± 0.02 cm (p = 6.75e-08) and in the *UBTF*-mutant NPCs it was 0.09 ± 0.03 cm compared to control NPCs 0.05 ± 0.01 cm (p=0.01), as determined by Wilcoxon signed-rank test. Example images of control (b) and patient (c) are shown in Fig 5b-c.

**Fig 5.**
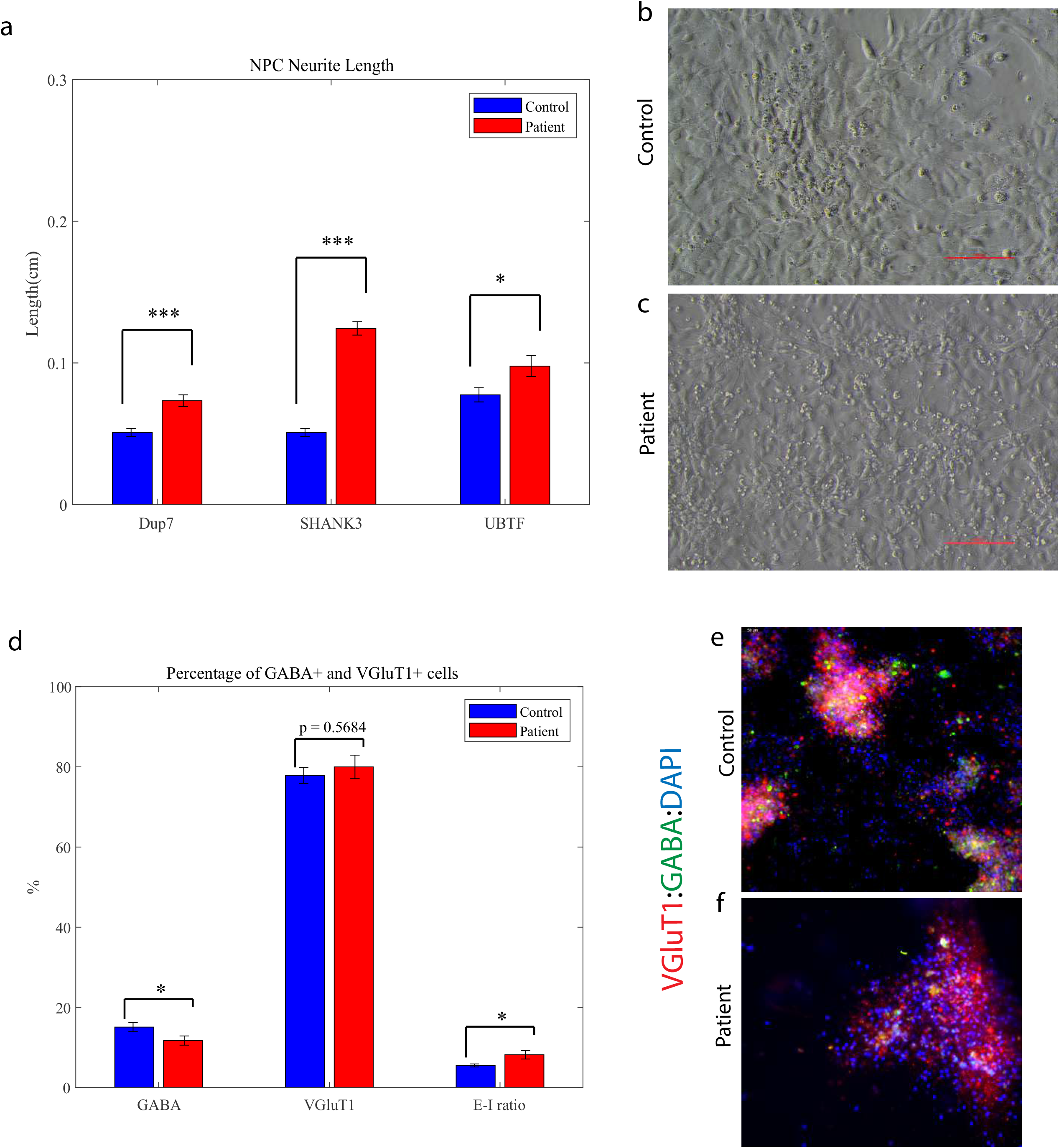
Mature young ASD-related mutant NPCs and an aberrant Excitation-Inhibition ratio in mutant neurons compared to controls. **a** A longer neurites in Dup7-NPCs (left) p=4.36e-04, *SHANK3*-NPCs (middle) p = 6.75e-08 and *UBTF*-NPCs (right) p= 0.01 mutants compared to control groups. **b** The averages of the percentage of GABA-expressing neurons (left) out of the total neurons in the *ASD*-mutant neuronal cultures are significantly lower than the percentage of GABA-expressing neurons in the control neuronal cultures (p=0.04), The averages of the percentage of VGluT1-expressing neurons (middle) out of the total neurons in the *ASD*-mutant neuronal cultures are similar to the percentage of VGluT1-expressing neurons in the control neuronal cultures (p=0.56) and a higher E-I ratio (right) in *ASD*-mutant neuronal cultures compared to control cultures (p = 0.03). **c**,**d** A representative image of mutant (**c**) and control (**d**) neurons that were immunostained for DAPI, GABA, and VGluT1.

Next, we used *GABA* and *VGluT1* antibodies to confirm the existence of GABA and *VGluT1*-positive cells in our cultures. Indeed, the control cultures contained 15.08 ±1.13 % *GABA*-expressing and 77.87± 2% *VGluT1*-expressing neurons. However, the ASD-related mutant cultures contained only 11.73±1.14 % *GABA*-expressing and 79.99 ± 2.93% VGluT1-expressing neurons (P_*GABA*_ = 0.04, P_*VGluT1*_ = 0.56); Additionally, we observed a higher ratio of Excitation-Inhibition (E-I ratio) in the Patient group compared to Control (8.183 ±1.05 and 5.51 ± 0.38 respectively;p=0.03) as shown in Fig 5d. Fig 5e-f shows an example image of control (e.) and mutant (f) neuronal cultures that were immunostained for DAPI (blue), *VGluT1* (red), and *GABA* (green).

## DISCUSSION

In this study, iPSC technology was used to investigate the physiological features of cortical neurons derived from human patients with different ASD-related mutations: *Dup7, GRIN2B, SHANK3*, and *UBTF*. For that purpose, we differentiated patient-derived iPSCs into cortical neurons^55^ since cortical alterations and malformations within the brain were identified in these mentioned mutations^14,29,43,50^.

A broad range of mutations has been associated with ASD, sharing autistic-like behaviors such as difficulties in social communication and interaction and restricted or repetitive behaviors or interests. The broad spectrum of mutations is characterized by affecting different genes and pathways. Various genes affecting the cytoskeletal^58–62^ and microtubule^63^ dynamics are altered in ASD-associated genes, disturbing a critical stage in axons and dendrites development^64^; For *Dup7* mutation, for example, a rare genetic syndrome caused by a micro-duplication in section q11.23 of chromosome 7, alterations in the ELN gene such as inherited and *de novo* deletions were reported, coding for the extracellular matrix protein, elastin, associated with connective-tissue malformations as reported in human patients^12^. *SHANK3* is a gene located at the terminal long arm of chromosome 22, coding for a master scaffolding protein found in the body’s tissues, and importantly in the brain, and has a critical role in the postsynaptic density of glutamatergic synapses and synaptic functions in rat and human brains^15,19^. The *GRIN2B* mutation results in a production of a nonfunctional GluN2B protein. A shortage or dysfunction of this protein may cause an extreme reduction in the number of the functional NMDA receptors^35^ causing neuronal impairments in both mice and human models^65,66^. *UBTF* is a gene coding for UBF, a transcription factor in RNA Pol I, critical for rRNA transcripts synthesis from rDNA in the nucleolus^45^. Loss of UBF induces nuclear disruptions, including inhibition of cell proliferation, rapid and synchronous apoptosis, and cell death in mice models^46^.

Although these mutations are functionally very different from one another, they cause similar symptoms in the patients. Interestingly, we observed a hyperexcitability pattern in all these mentioned ASD-related mutations in an early-stage cell development (3-5 weeks post differentiation) that we measured by electrophysiological recordings. This hyperexcitability involved different aspects; In terms of sodium-potassium currents and activity, a consistent increase in sodium currents was observed within the four patient-derived neurons, which could, in turn, increase the excitability of the neurons by decreasing the action potential threshold^67^. These alterations of sodium currents can lead to abnormal neuronal activity, a phenomenon that also occurs in epilepsy^68^. It is interesting to note that all these four mutations have a strong association with epilepsy, and many of the patients also suffer from epilepsy^14,27,44,53^. We also observed an increase in the EPSC rate and amplitude, indicating pre and postsynaptic changes that occur in the mutant neurons compared to the controls neurons. Similar findings were reported in mice models of autism ^69–73^. Furthermore, we observed more evoked action potentials in response to current stimulation in the mutant-derived neurons, alongside with a decrease in *GABA*-positive cells and an increase in the E-I ratio, which may contribute to the occurrence of infantile epileptic seizures in affected human subjects. We previously reported such changes also in another ASD model of an *IQSEC2* mutation^74^. Furthermore, the NPCs derived from individuals with ASD-related mutations display longer neurite-like branches compared to control groups supporting our conclusions of their early maturation.

All these changes can indicate that the ASD mutant neurons develop faster, and at this early stage, when the control neurons are still very immature, they are already spiking and connecting with other neurons.

Several previous genetic studies showed a rise in cortical activity by documenting excitation-to-inhibition ratio system alterations^74–76^, suggesting that periodic seizures and sensory hyperreactivity in ASD are caused by cortical hyperexcitability^77,78^, which was also previously observed in fragile-X syndrome^79^. Previously, we reported a similar early time point hyperexcitability pattern in another ASD-related mutation -the A350V IQSEC2mutation^74^. In that study, we followed the IQSEC2-mutant neurons that started more active and more connected (5 weeks post differentiation) as they became hypoexcitable with reduced synaptic connections later on in the differentiation process. An additional hyperexcitability pattern in 15q11-q13 Duplication Syndrome, a different ASD-associated mutation, was reported^80^. A reduction in synaptic connections was reported in a long line of ASD-related and ID-related studies using mice and human models^81–84^, suggesting a potential shared accelerated aging mechanisms in ASD-associated mutations^62,63,85–87^.

We speculate that there may be a connection between this early hyperexcitability and the later synaptic deficits, as perhaps this early hyperexcitability is neurotoxic to the cell at such an early stage of development.

Since, especially with mice studies, it is much harder to measure this early developmental stage, perhaps this is a stage that precedes the synaptic degradation in many other ASD mutations. More evidence of this early maturation was presented in a study with ASD patients with macrocephaly where the neurons derived from the patients were more arborized early in the development, and gene expression profiles also suggested an earlier maturation^88^. More evidence for functional hyperactivity in epilepsy and ASD-related mutations in human models were reported; briefly, an engineered iPSC-derived neuron with the homozygous P924L mutation (one of many epilepsy-associated Slack mutations) displayed increased K_Na_ currents and more evoked action potentials in both single neurons and a connected neuronal network^89^.

Our Findings present a shared phenotype of early maturation and hyperexcitability in four ASD-related mutations using patient-derived cortical neurons, indicating that there may be a common neurophysiological phenotype in ASD-related variants, sharing similar behavioral phenotypes but a different genotype. iPSC-derived neurons were previously used as a research tool for investigating physiological and cellular alterations characterizing various disorders including autism^74,88,90,91^,epilepsy^89,92,93^ and other neurodegenerative diseases such as Parkinson’s^94–97^ and schizophrenia^98–100^. Here we concentrated on the early developmental physiological alterations in 4 different ASD and epilepsy-related genes. The enhanced maturation and excitability in such young neurons may be deleterious to the cells and may later result in synaptic degeneration as was previously described in neurons derived from ASD and epilepsy patients^74,84,88–90,92,93^.

## Acknowledgments

This work was supported by Zuckerman STEM Leadership Program, Israel science foundation (ISF) grants 1994/21 and 352/21, new childhood – The organization for *UBTF*, Almy Foundation, and the *GRIN2B* Disorder Research Foundation (GDRF). We would like to thank the Technion Genomics Center for their contribution to STR testing services.

---

**Fig S1. STR and PCR tests validate the existence of the mutations in Dup7 and *SHANK3***.

**a** The sample profiles show that the PBMC and NPC lines of UOHi011 completely match one another. **b-c** PCR validation of the (C.3679insG) *shank3* mutation in the mutant line and not in the control line.

