## Supplementary Fig. S1 for "Early maturation and hyperexcitability is a shared phenotype of cortical neurons derived from different ASD-associated mutations"

a

Cell lines' ID: UOHi011_O_Dup7_NPC, UOHp011 _Dup7_PBMCs

| Marker | UOHi011_O_Dup7_NPC | UOHp011_Dup7_PBMCs |
|---|---|---|
| AMEL | X,Y | X,Y |
| D3S1358 | 16,18 | 16,18 |
| D1S1656 | 15,17 | 15,17 |
| D2S441 | 11 | 11 |
| D10S1248 | 15,16 | 15,16 |
| D13S317 | 11,12 | 11,12 |
| Penta E | 7,10 | 7,10 |
| D16S539 | 9,11 | 9,11 |
| D18S51 | 13,15 | 13,15 |
| D2S1338 | 20,24 | 20,24 |
| CSF1PO | 11,12 | 11,12 |
| Penta D | 10,11 | 10,11 |
| TH01 | 6,9.3 | 6,9.3 |
| vWA | 17 | 17 |
| D21S11 | 28,32.2 | 28,32.2 |
| D7S820 | 9,10 | 9,10 |
| D5S818 | 11 | 11 |
| TPOX | 8 | 8 |
| DYS391 | 10 | 10 |
| D8S1179 | 14,16 | 14,16 |
| D12S391 | 19,22 | 19,22 |
| D19S433 | 13.2,15 | 13.2,15 |
| FGA | 22 | 22 |
| D22S1045 | 15,16 | 15,16 |

**Summary**

The samples profiles show that lines UOHi011_O_Dup7_NPC and UOHp011 _Dup7_PBMCs completely match one another.

b

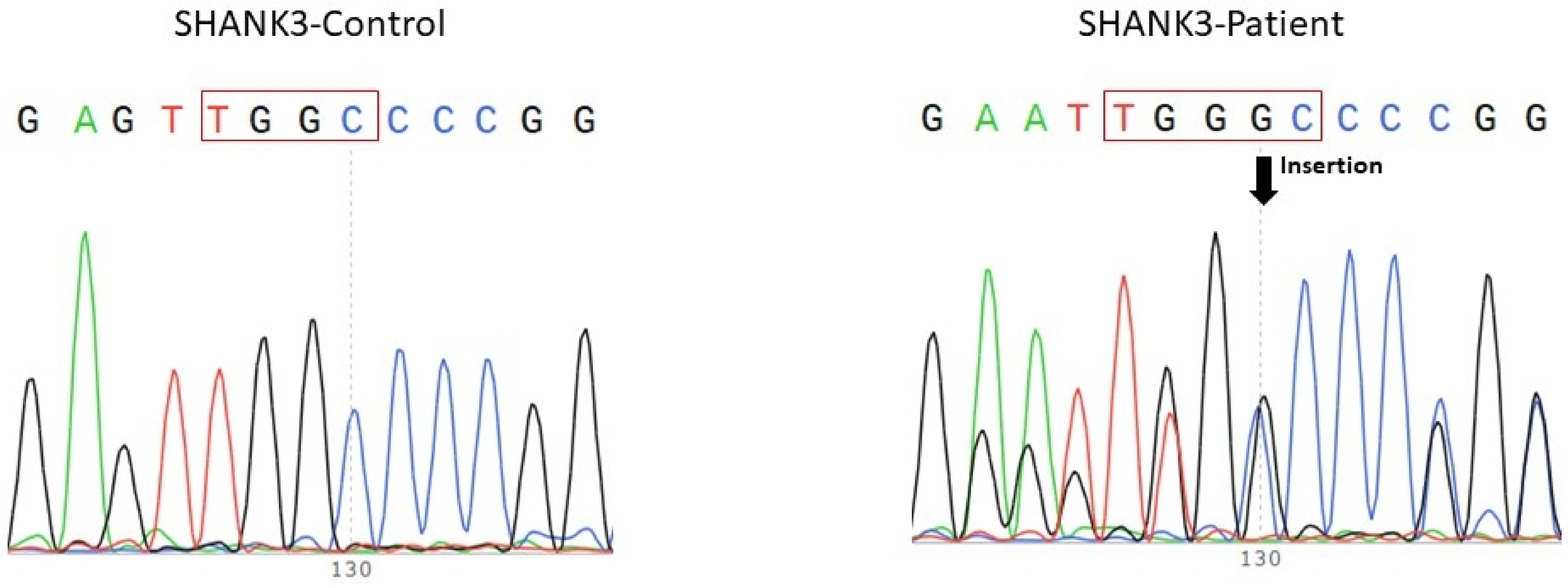
