## Supplementary Table S1 for "Early maturation and hyperexcitability is a shared phenotype of cortical neurons derived from different ASD-associated mutations"

**Table T1.** A description of the patients' cohort

| **Patient ID** | **Mutation** | **Sex** | **Age at sampling (Y)** | **Clinical symptoms** |
| --- | --- | --- | --- | --- |
| UOHi011 | Dup7 (7q11.23 dup) | M | 7.5 | ASD, minor ID |
| UOHi010 | Control | M | 39 | - |
| 784313 | GRIN2B (c.2065 G->T) | F | 2 | ASD, ID |
| UOHi010 | Control | M | 39 | - |
| UOHi003-A | SHANK3 (C.3679insG) | F | 10 | ASD, ID |
| UOHi002-A | Control | F | 47 | - |
| UOHi004 | UBTF (E210K) | F | 22 | ASD, ID |
| UOHi005 | Control | F | 50 | - |
