## Supplementary Table S2 for "Early maturation and hyperexcitability is a shared phenotype of cortical neurons derived from different ASD-associated mutations"

**Table T2.** Spike shape analysis (non-parametric statistical tests: Wilcoxon signed rank test) of mutant and control cortical neurons.

|  | **Dup7** | **GRIN2B** | **SHANK3** | **UBTF** |
| --- | --- | --- | --- | --- |
| **Spike Threshold** | µ_mutant_ =27.6 ± 6.5 mV  µ_Ctl_ =21.09 ± 12.05 mV | µ_mutant_=21.5 ± 3.6 mV  µ_Ctl_=22.2 ± 6.7 mV | µ_mutant_=30.4 ± 5.6 mV  µ_Ctl_=25.5 ± 4.2 mV | µ_mutant_=32.1 ± 4.9 mV  µ_Ctl_=21.5 ± 16.7 mV |
|  | P = 0.29 | P = 0.27 | **P = 0.02 *** | P = 0.07 |
| **Spike Height** | µ_mutant_=25.4 ± 15.5 mV  µ_Ctl_=27.6 ± 19.4 mV | µ_mutant_=41.2 ± 12.5 mV  µ_Ctl_=20.5 ± 14.01 mV | µ_mutant_=44.3 ± 11.4 mV  µ_Ctl_=34.4 ± 7.5 mV | µ_mutant_=50 ± 17.05mV  µ_Ctl_=36.04 ± 18.01 mV |
|  | P = 0.14 | **P = 0.008 **** | P = 0.09 | **P = 0.03 *** |
| **Spike width** | µ_mutant_ =3.2 ± 3.4 ms  µ_Ctl_ =2.4 ± 3.4 ms | µ_mutant_=8.3 ± 5.8 ms  µ_Ctl_=27.6 ± 19.4 ms | µ_mutant_=6.5 ± 5.9 ms  µ_Ctl_=5.9 ± 1.2 ms | µ_mutant_=3.3 ± 1.5 ms  µ_Ctl_=12.5 ± 17.8 ms |
|  | P = 0.7 | P = 0.065 | P = 0.17 | **P = 0.003 **** |
| **Rise time** | µ_mutant_ =1.9 ± 3.1 ms  µ_Ctl_ =0.68 ± 0.87 ms | µ_mutant_=7.2 ± 5.1 ms  µ_Ctl_=3.6 ± 1.9 ms | µ_mutant_=4.6 ± 2.6 ms  µ_Ctl_=4.6 ± 1.8 ms | µ_mutant_=2.5 ± 1.6 ms  µ_Ctl_=5.2 ± 5.3 ms |
|  | P = 0.34 | P = 0.23 | P = 0.58 | P = 0.18 |

*P<0.05, **P<0.01, ***P<0.001 and ****P<0.0001
